# SigDyn: single-cell mutational signature dynamics

**DOI:** 10.64898/2026.01.20.700537

**Authors:** Sara B. Costa, Joana P. Gonçalves

## Abstract

**Motivation:** Cancer is driven by mutations that confer tumor cells proliferation advantages but also vulnerabilities with therapeutic potential. Underlying tumor progression are mutational processes that imprint signatures of co-occurring mutation types in tumor cell genomes. The discovery of signatures and their etiologies offers insight into disease mechanisms and treatment opportunities, however acquisition of mutations by tumor cells over time leads to intratumor heterogeneity that may challenge treatment efficacy. Understanding which mutational processes contribute to the evolution of each tumor can reveal therapeutic strategies, but it requires single-cell mutational signature analysis, which remains unexplored.

**Results:** We present SigDyn, a framework enabling analysis of tumor mutational signature dynamics at single-cell resolution. To do this, SigDyn combines variant detection, tumor phylogeny inference, and signature identification. We applied SigDyn to scRNA-seq of invasive ductal carcinoma (IDC) and laryngeal squamous cell carcinoma (LSCC). Inferred phylogenies recapitulated tumor markers and progression from normal tissue to tumor metastasis. Tumor-specific processes beyond aging, like cellular disruptions or patient treatments, were only exposed at single-cell level reinforcing the importance of granularity. For IDC, prominence of signature SBS26 with tumor evolution pointed to mismatch repair (MMR) deficiency, supported by downregulation of MMR genes. For LSCC, metastasis enrichment with SBS32-active cells and immune markers suggested influence of immunosuppressant treatment azathioprine, linked to increased risk of SCCs. The dynamics uncovered by SigDyn confirm its potential as a tool to investigate mechanisms of tumor evolution and heterogeneity.

**Availability:** github.com/joanagoncalveslab/sigdyn.

**Contact:**. Supplementary Information: included.

## Introduction

Cancer arises due to a combination of driver mutations that disrupt cellular growth and death, enabling cells to proliferate uncontrollably. The progression of a tumor is further fueled by an accumulation of mutations that leads to heterogeneity in the genetic landscape of tumor cells (Fidler, 1978), posing critical challenges to treatment efficacy (McGranahan and Swanton, 2017). Understanding tumor evolution at single-cell level is thus important to elucidate mechanisms driving tumor progression and discover strategies to overcome treatment resistance.

Existing computational approaches to analyze tumor evolution and heterogeneity fit into two main categories. The first studies genomic variation events directly, while the second aims to capture signatures left by the mutational processes underlying the mutations acquired during tumor progression.

Numerous methods have been proposed to characterize the diversity of cell populations within a tumor focusing on genomic variation events. Advances in sequencing technology have enabled the desired resolution to study such events at single-cell level, with scRNA-seq data being particularly appealing for its wide availability and cost-effectiveness compared to scDNA-seq. Most approaches capture heterogeneity through subclone identification and/or uncover the sequence of variation events underlying tumor evolution using phylogeny inference, based on either copy number variants (CNVs, Tirosh *et al*. (2016); Gao *et al*. (2021)) or single-nucleotide variants (SNVs, Jahn *et al*. (2016); Aaltonen *et al*. (2020); Marot-Lassauzaie *et al*. (2024)). Both have had success but also harbor limitations: CNV-based methods assume that observed changes in coverage or gene upregulation emerge from genomic CNV, overlooking possible transcriptional activation; SNV-based methods face challenges in variant detection from individual cells, due to low molecular input. Alternative methods use genetically-engineered lineage tracing systems to facilitate reconstruction of tumor evolution (Quinn *et al*., 2021; Yang *et al*., 2022).

The second category of methods focuses on processes responsible for mutations acquired during tumor progression, which are known to leave distinctive mutational signatures of co-occurring mutation types in the genomes of tumor cells (Alexandrov *et al*., 2013b). Various signatures have been associated with relevant mutagenic processes, including exposure to exogenous mutagens such as tobacco smoking, or deficiencies in cellular mechanisms like DNA repair (Alexandrov *et al*., 2013b). Signatures linked to defective homology-directed repair (HDR) have specifically been reported to identify breast tumors with functional BRCA1/2 deficiency, indicated for PARP inhibitor therapy, which had no detectable mutations in those HDR genes (Davies *et al*., 2017). The therapeutic significance of mutational signatures has inspired the development of approaches that analyze signature activities over a pseudo-timeline of evolution inferred by clustering variant allele frequencies from bulk genome sequencing of a tumor (Rubanova *et al*., 2020), possibly in combination with the detection of tumor subclones (Abécassis *et al*., 2021). Nevertheless, such methods may be limited in their ability to distinguish evolution branches based on bulk aggregates of mixed cell populations, which could mask subclonal mutational dynamics.

Tracing tumor mutational processes at single-cell level can reveal therapeutic opportunities that might be missed by bulk analyses, and is yet to be explored. We argue this is a promising avenue, since many studies have successfully characterized single-cell genomic variation. In addition, mutational signatures look at changes in sequence, aggregated over an entire genome and agnostic to genomic location, making signature detection relatively robust to individual mutations.

We present SigDyn, a framework enabling the analysis of mutational signature dynamics of a tumor at single-cell resolution. To achieve this, SigDyn exploits both genomic variation events and their underlying generative mutational processes, in three steps: (1) detection of somatic variants from tumor cells, (2) inference of a tumor evolution trajectory, and (3) estimation of mutational signature dynamics over the inferred tumor evolution. To showcase the capabilities of SigDyn, we assess each of its methodological components and investigate its application to scRNA-seq data of two tumors: an invasive ductal carcinoma and a laryngeal squamous cell carcinoma.

## Materials and methods

The SigDyn method reveals mutational signature dynamics from single-cell sequencing of a tumor sample in three steps (Fig. 1). First, it identifies somatic variants across the tumor cells. Second, it infers a tumor evolution trajectory by reconstructing a phylogenetic tree based on the tumor cell genotypes. Third, it estimates mutational signature activities for tumor cells and inferred ancestor nodes in the tree, to characterize signature dynamics over the inferred tumor evolution.

**Figure 1.**
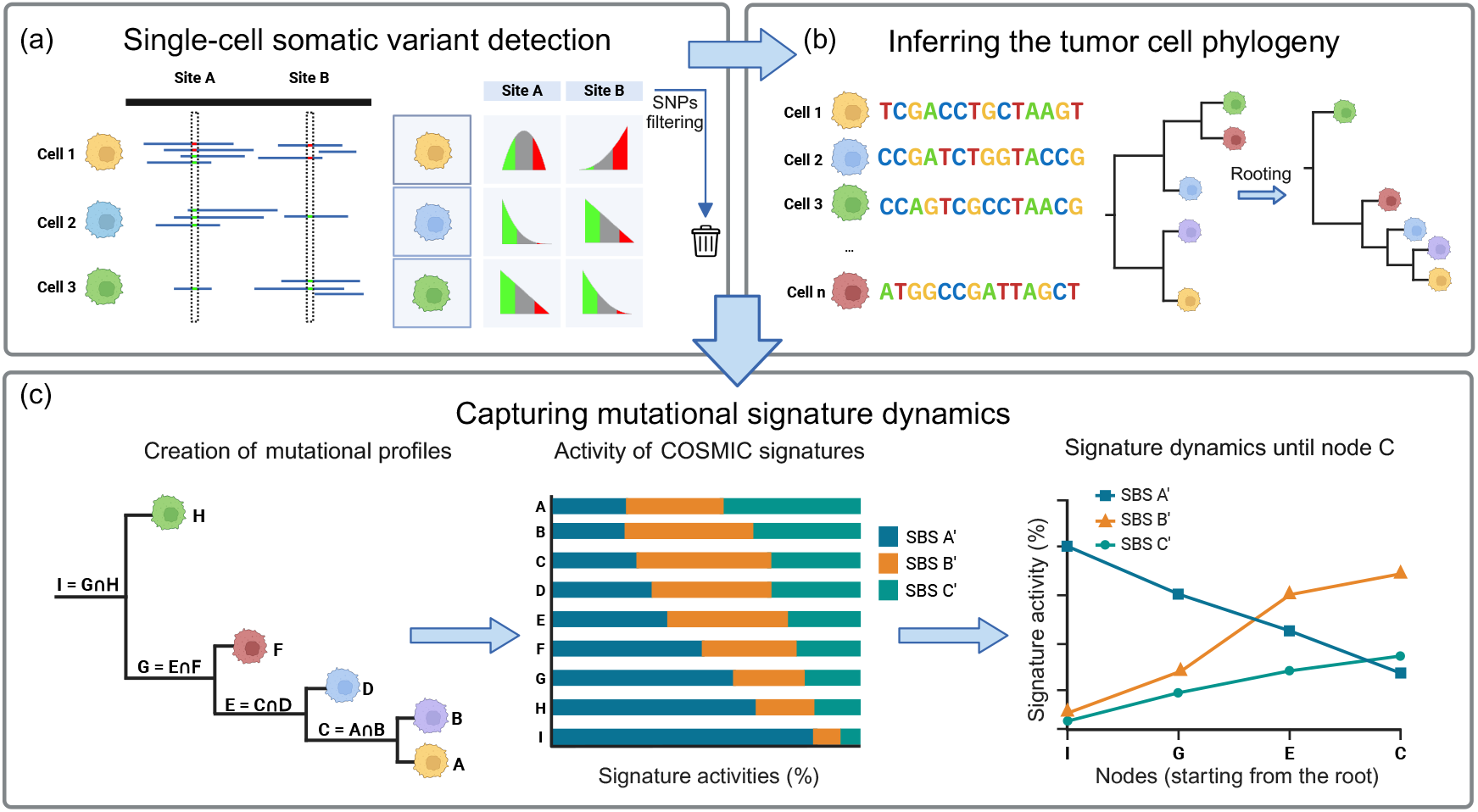
SigDyn framework. (a) Variants are detected from single-cell RNA-seq data across individual cells. Two example variant sites, with red and green indicating alternate and reference alleles, respectively. Using beta-binomial distributions, genotypes are assigned based on the highest probability across three categories: green, homozygous reference; gray: heterozygous; red: homozygous alternate. Finally, variants found in a public Panel of Normals (PON) are filtered to remove common artifacts and germline variants. (b) Phylogeny inference to capture evolutionary relations between cells. Tree rooted using the cell with the largest number of reference alleles, establishing a baseline for mutation accumulation over the tree. (c) Ancestral node profiles are created by aggregating common mutations from children nodes, and mutational signatures are identified for single-cell and ancestor profiles to trace signature dynamics over the tumor evolution.

### Step 1: Single-cell somatic variant detection

The first step of SigDyn identifies somatic variants in tumor cells. This is performed using the PhylinSic pipeline (Liu *et al*., 2023), which combines probabilistic modeling with local smoothing and imputation to mitigate the sparsity of single-cell data when calling variants for individual tumor cells. The variants obtained are further processed by SigDyn to filter germline polymorphisms and sequencing artifacts, aiming to retain somatic events. The final set of variants is used to obtain the genotype of each tumor cell.

Briefly, PhylinSic first identifies candidate sites showing variation across the tumor cells based on a pseudo-bulk aggregate of the reads of all individual cells. Candidate sites are then analyzed to call variants at the single-cell level. For each site, PhylinSic models the observed allele frequencies using a beta-binomial distribution that seeks to capture both biological variation and technical noise in sequencing data.

Based on the probability distribution, PhylinSic assigns one of three states: homozygous reference (RR), heterozygous (RA), or homozygous alternate (AA), according to the respective interval of allele frequency with the highest probability mass. To mitigate sparsity, PhylinSic applies k-nearest neighbor (kNN) smoothing, leveraging similarities between cells to impute uncertain calls and refine assignments. PhylinSic is used with default parameters.

After variant calling, SigDyn additionally filters sites likely denoting germline variants. When matched normals are available, sites classified as homozygous alternate (AA) in cells from both tumor and normal are considered germline and removed if detected in over 40% of the cells in the normal sample. In the absence of matched normals, a Panel of Normals (PON) from GATK (Poplin *et al*., 2018) is used to exclude all PON sites, and thus filter sequencing artifacts and common polymorphisms identified across a cohort of 1000 human genomes (Auton *et al*., 2015).

### Step 2: Inferring a tumor cell phylogeny

The second step of SigDyn infers a phylogeny to capture evolutionary relations between the tumor cells and estimate the order by which they likely emerged during tumor progression.

The phylogeny is inferred based on the genotypes of tumor cells, represented as nucleotide sequences, with each position denoting one of the sites identified as varying across the cells in Step 1. Following Liu *et al*. (2023), homozygous reference and variant states are represented by their respective nucleotides, whilst heterozygous states are assigned a randomly chosen nucleotide, different from the reference and the variant alleles. Heterozygous states are therefore consistently represented by the same nucleotide across all cells with a heterozygous call for the same position. All genotypes are of equal length and order, ensuring each position in the sequence corresponds to the same genomic site across all cells.

To infer the tumor cell phylogeny, SigDyn uses FastTree 2 (v2.1.11) (Price *et al*., 2010), using the default Jukes–Cantor model of nucleotide evolution. We selected FastTree 2 because this maximum-likelihood approach allows us to make fewer assumptions and rely mainly on sequence similarity, without incorporating more complex evolutionary priors that might not fit the underlying tumor evolution. This is also particularly suitable given that the genotypes contain variant sites only and are not full-length genomic sequences. In addition, FastTree 2 scales well to large datasets, supporting wider applicability.

To provide guidance on the direction of tumor evolution, SigDyn allows to root the tree using a reference or baseline genotype. In practice, this could be based on a cell selected from a matched normal sample, if available. Otherwise, SigDyn roots the tree based on the cell with the largest count of reference (RR) sites, or in other words the least mutated cell. This aligns with the goal of modeling tumor evolution, with mutations expected to accumulate from the root to the leaves. Rooting is performed using FigTree (v1.4.4) (FigTree, 2024).

### Step 3: Capturing mutational signature dynamics

The final step of SigDyn identifies mutational signatures active in the tumor cells. Signature activities are integrated with the tumor phylogeny inferred in Step 2 to enable analysis of single-cell mutational signature dynamics underlying tumor evolution.

Mutational signatures can be identified de novo, or estimated together with their activities based on large collections of signatures from relevant databases. For clarity, the activity of a signature denotes the number of mutations contributed by such signature to the mutational profile of a cell. Here, we estimate activities for mutational signatures available in the Catalogue of Somatic Mutations in Cancer (COSMIC v3.4, Tate *et al*. (2019)), which have been originally identified from whole-genomes and whole-exomes of patient tumors using the SigProfiler non-negative matrix factorization approach (Alexandrov *et al*., 2013a). Specifically, we focus on the 86 COSMIC mutational signatures described over 96 possible types of single-base substitutions within a trinucleotide context (96 = 6 substitution types *×* 4 possible 5’ neighbor bases *×* 4 possible 3’ neighbor bases).

To enable the analysis of signature activities over the entire phylogenetic tree, we first create the mutational profile of each tumor cell and each ancestor tree node (Fig. 1c, left), based on which signatures can be identified. For a tumor cell, the mutation profile directly tallies the somatic variants identified in Step 1 corresponding to each of the 96 possible types of single-base substitutions. For an ancestor node, the mutational profile is created in a similar way, but then based on the set of somatic variants shared by all the children of that node. This follows from the infinite sites assumption, whereby an acquired mutation will become fixed and will be propagated to all descendants, meaning that a parent node should carry the intersection of its children’s variant sets (containing the variants present before the tree split).

Signature activities are obtained for each tumor cell or ancestor node using non-negative least squares (NNLS, Fig. 1c, middle), to estimate the combination of signatures and activities that maximize the reconstruction of the mutational profile of the cell or ancestor node. We initially fit all 86 COSMIC signatures using SigProfilerAssignment (Díaz-Gay *et al*., 2023) to the profiles of tumor cells, to avoid premature exclusion of potentially relevant processes. We then retained only those signatures present in at least 10% of tumor cells, which were designated as active signatures. We further refitted only the active signatures to both tumor cells and internal ancestor nodes using NNLS. This was done to mitigate artifacts that could arise from fitting marginally detected mutational signatures. With mutational signature activities assigned to every node in the phylogeny, SigDyn captures how signature activities change from the root to any node or cell of interest (Fig. 1c, right).

### Case Studies

#### Data and preprocessing

We applied SigDyn to scRNA-seq data from two tumor samples, both obtained using droplet-based 10x Genomics technology. The first sample was a biopsy of an invasive ductal carcinoma (IDC) from a 65-year-old female patient (10x Genomics, 2021). We used available pre-aligned BAM files and gene count matrices, originally generated using Cell Ranger v6.0.0. The second sample contained cells collected from 4 regions of a laryngeal squamous cell carcinoma (LSCC) from a 60 to 65 year-old male patient (patient 3, Sun *et al*. (2024)): normal laryngeal mucosa (N), tumor margins (R), carcinoma *in situ* (T), and lymph nodes metastasis (L). Reads from each region were aligned to the human reference genome (hg38) using Cell Ranger v8.0.1 to generate both alignment BAM files and gene expression counts.

#### Evaluation and analysis of SigDyn results

##### Tumor evolution

To assess the SigDyn-inferred phylogeny, we compared FastTree 2 to an alternative Bayesian inference method BEAST 2, used by PhylinSic (Liu *et al*., 2023). We set BEAST 2 parameters as in Liu *et al*. (2023), using a relaxed log-normal clock (RLN), a generalized time-reversible site model (GTR) of nucleotide substitution, and Yule tree branching priors. We also followed the convergence assessment procedure of visualizing the posterior tree distribution with a DensiTree plot and confirming that the posterior probability trace reached a stable plateau. To further investigate how the inferred phylogeny captured genotypic relations between single-cells, we analyzed phylogenetic distance and genotype similarity of all pairs of tumor cells. Phylogeny-based distance was defined as the sum of branch lengths along the unique path connecting the respective leaves of the pair of tumor cells in the phylogenetic tree, up to their closest ancestor. Genotype similarity was calculated as the proportion of genotype positions showing matching nucleotides for both tumor cells.

##### Tumor clones and regions

We identified IDC clones to enable the analysis of gene expression changes and mutational signature dynamics along the inferred tumor phylogeny. To obtain the clones, we applied centroid-linkage hierarchical clustering to the pairwise distances between the tumor cells in the phylogenetic tree, and used a distance threshold of 0.9 to define the clones. For LSCC, we analyzed gene expression and signature dynamics over the 4 tumor regions originally available.

##### Gene expression analysis

Gene counts were normalized to a total of 10^4^ per cell to mitigate library-size differences, followed by log-transformation to stabilize the variance across genes, according to best practices for scRNA-seq (Heumos *et al*., 2023). For dimension reduction analysis of tumor cell expression patterns, gene expression was further standardized to zero mean and unit variance, the top 2,000 most variable genes were selected, and UMAP was applied as implemented in Scanpy (Wolf *et al*., 2018).

For differential gene expression analysis (DGEA) between cell groups of interest, such as cells with a given active or inactive mutational signature, we used MAST (Finak *et al*., 2015): a two-part generalized linear model designed for zero-inflated scRNA-seq data. As input to MAST, we provided the normalized and log-transformed expression values of genes expressed in at least 10% of the cells, again following recommended practice (Heumos *et al*., 2023). We used MAST to contrast the two cell groups of interest, and corrected the resulting *p*-values from the likelihood ratio tests for multiple testing using the Benjamini-Hochberg method. Genes were considered differentially expressed when *p*_adj_ *<* 0.01 and | log_2_(FC)| *>* 0.5. To identify relevant biological processes annotated with the differentially expressed genes, we performed overrepresentation analysis separately for up- and down-regulated genes against the GO Human Biological Process 2021 database with the enrichr tool, using the gseapy package (Fang *et al*., 2023).

## Results

We investigated the potential of SigDyn in two ways. First, we examined the performance of each methodological component of SigDyn separately using the smaller breast IDC dataset, focusing on: somatic variant detection, tumor evolution reconstruction, and mutational signature identification. Second, we analyzed the mutational signature dynamics obtained using SigDyn for the breast IDC and the head-and-neck LSCC datasets.

### Somatic variant detection

The first step of SigDyn identifies somatic variants for each single-cell in a given tumor sample. For the breast IDC dataset, SigDyn detected 776 somatic variants across the 687 cells, following single-cell variant calling with PhylinSic and further filtering of germline polymorphisms based on the 1000 Genomes panel of normals (see Methods). Of those 776 variants originally called, 46% were categorized as homozygous alternate (AA) mutations in at least one cell, with an average incidence of 22% per single-cell (Fig. 2a). Overall, 41% of variants (316/776) were filtered, 81% of which were categorized as AA in at least one cell (Supplementary Fig. S1). Notably, the fact that most AA variants were filtered as likely germline (255*/*356 ≈ 72%, Supplementary Fig. S1) underlined the importance of the additional filtering performed by SigDyn to improve the detection of tumor somatic mutations, which SigDyn can use to characterize tumor evolution via phylogeny and mutational signature analysis.

**Figure 2.**
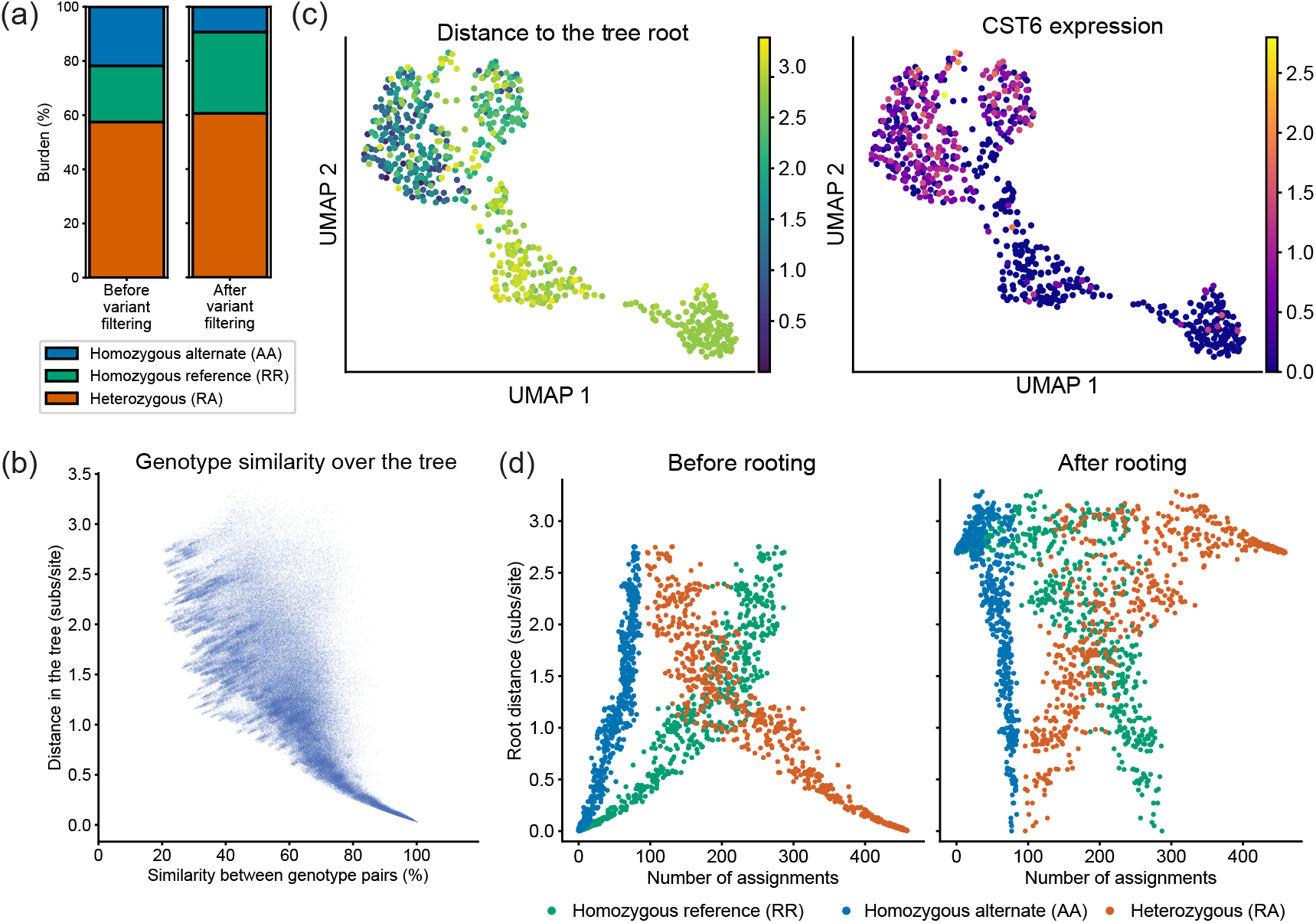
Somatic variants, single-cell genotypes, and tumor cell phylogeny inferred by SigDyn for the breast IDC sample: (a) proportion of variant sites (left) identified using PhylinSic, and (right) upon filtering by SigDyn based on the 1000 Genomes panel of normals, stratified per category; (b) relation between single-cell genotype similarity and distance in the tumor phylogenetic tree inferred by SigDyn using FastTree 2; (c) left, relation between phylogeny and single-cell gene expression (UMAP embedding); (c) right, expression of CST6 tumor suppressor gene; and (d) effect of explicit tree rooting using the cell with the lowest somatic variant count on the relation between the distance of each cell to the root of the phylogenetic tree and the number of variants identified for such cell.

### Tumor evolution reconstruction

The second step of SigDyn infers a phylogenetic tree to capture evolutionary relations between tumor cells based on genotype similarity. Here, SigDyn uses FastTree 2 (Price *et al*., 2010), a maximum-likelihood algorithm relying on a simple Jukes-Cantor model of nucleotide evolution. This approach was chosen for: making few assumptions about sequence changes shaped by unique dynamics and selection pressures in tumor evolution; scaling to large data, thus being well suited for scRNA-seq.

We assessed the breast IDC phylogeny inferred by FastTree 2, and compared it to the BEAST 2 (Bouckaert *et al*., 2014) Bayesian approach in PhylinSic (Liu *et al*., 2023). FastTree 2 inference completed within a few seconds whereas BEAST 2 needed a considerable number of iterations (Supplementary Fig. S2), and in excess of hours or days to converge.

#### Cell sequence similarity translates to phylogenetic distance

The phylogenies inferred by SigDyn using FastTree 2 revealed a desired association between single-cell genotype similarity and phylogenetic distance (Fig. 2b), whereby cells with more similar genotypes were closer together in the tree and thus showed smaller phylogenetic distances. This relation was also apparent using the alternative BEAST 2 tree inference method, confirming that both approaches were able to capture it (Supplementary Fig. S2).

#### Phylogenetic structure aligns with gene expression patterns

Since the genotype of a cell can influence its cellular function, we expected that gene expression could show correspondence with the organization of cells in the phylogenetic tree. Notably, clones identified based on the SigDyn/FastTree 2 and BEAST 2 phylogenies were concordant with variation in gene expression, providing evidence of functional coherence in support of the phylogenetic relations (Supplementary Fig. S3). We further observed that expression of the tumor suppressor gene CST6 was anti-correlated with distance to the root in the SigDyn-inferred phylogeny (Spearman *ρ* = −0.51, *p*-value 1.71*×*10^−46^, Fig. 2c), confirming concordance between phylogeny and function for this regulator of cell proliferation involved in tumorigenesis (Jin *et al*., 2012; Rivenbark *et al*., 2006; Li *et al*., 2021).

#### Tree rooting provides a direction for tumor evolution

To guide the direction of tumor evolution, SigDyn can explicitly root the phylogenetic tree. Here we did so based on the cell with the largest number of reference sites, under the assumption that mutations accumulate over time as a tumor develops. To evaluate the effect of tree rooting, we compared the variant profiles of the cells in the tree inferred with and without this rooting (Fig. 2d). Regardless of rooting, in both trees, the numbers of homozygous reference (RR) and alternate (AA) sites per cell followed a similar trend, and both were anti-correlated with the number of heterozygous variants (RA). The increase in heterozygous (RA) sites was mostly mirrored by a decrease in homozygous reference (RR) sites, while the number of homozygous alternate (AA) sites remained more stable. Finally, the tree rooting achieved the desired effect, as cells in the explicitly rooted tree showed larger numbers of heterozygous (RA) sites with increasing distance from the root (Fig. 2d, right), while the opposite was observed for the tree without explicit rooting (Fig. 2d, left).

Our results show that FastTree 2 can efficiently infer a tumor phylogeny that captures the evolution of single-cell genotypes, concordant with gene expression changes. We highlight that SigDyn is modular and can be used with alternative phylogenetic inference methods, such as BEAST 2, whenever appropriate. Explicit tree rooting is optional. In general, we recommend rooting based on a reference genotype from a matched-normal sample of the same donor. If that is unavailable, rooting based on the cell with the lowest somatic variant count may be a reasonable compromise, given that tumor biopsies often also include normal tissue. However, this is not guaranteed and rooting based on a particular tumor cell could lead to suboptimal phylogeny inference if, for instance, different tumor lineages diverged early on and no shared ancestor is present in the data.

### Mutational signature identification

The final step of SigDyn estimates the activity of known tumor mutational signatures from the COSMIC database per single-cell in the IDC sample (Tate *et al*. (2019), Methods). To assess single-cell results, we also estimated activities for a pseudo-bulk aggregate of all somatic variants detected across the cells.

We detected 5 signatures at single-cell level, five in at least 10% of the cells: SBS1, SBS5, SBS12, SBS26, and SBS40c (Fig. 3a). Both single-cell and pseudo-bulk analyses identified mismatch repair deficiency (SBS26) and clock-like or aging (SBS1 and SBS40c) signatures (Nik-Zainal *et al*., 2012, 2016; Senkin *et al*., 2024). In single-cells, we detected another clock-like signature (SBS5), and also signature SBS12 of unknown etiology (Alexandrov *et al*., 2013b). Two pairs of signatures were highly similar, ranking second and fifth most similar among 3655 COSMIC signature pairs: SBS12/26 and SBS5/40c, with cosine similarities of 0.93 and 0.91 (Fig. 3b). The signatures in each pair were also complementarily detected, with one present and the other absent for most cells (Pearson’s correlation: SBS12 vs SBS26 -0.78, SBS5 vs. SBS40c -0.85), indicating they could not be easily distinguished. While we did not find literature relating SBS12 and SBS26, SBS5 and SBS40(c) have both been linked to polymerase zeta activity (Hwang *et al*., 2025).

**Figure 3.**
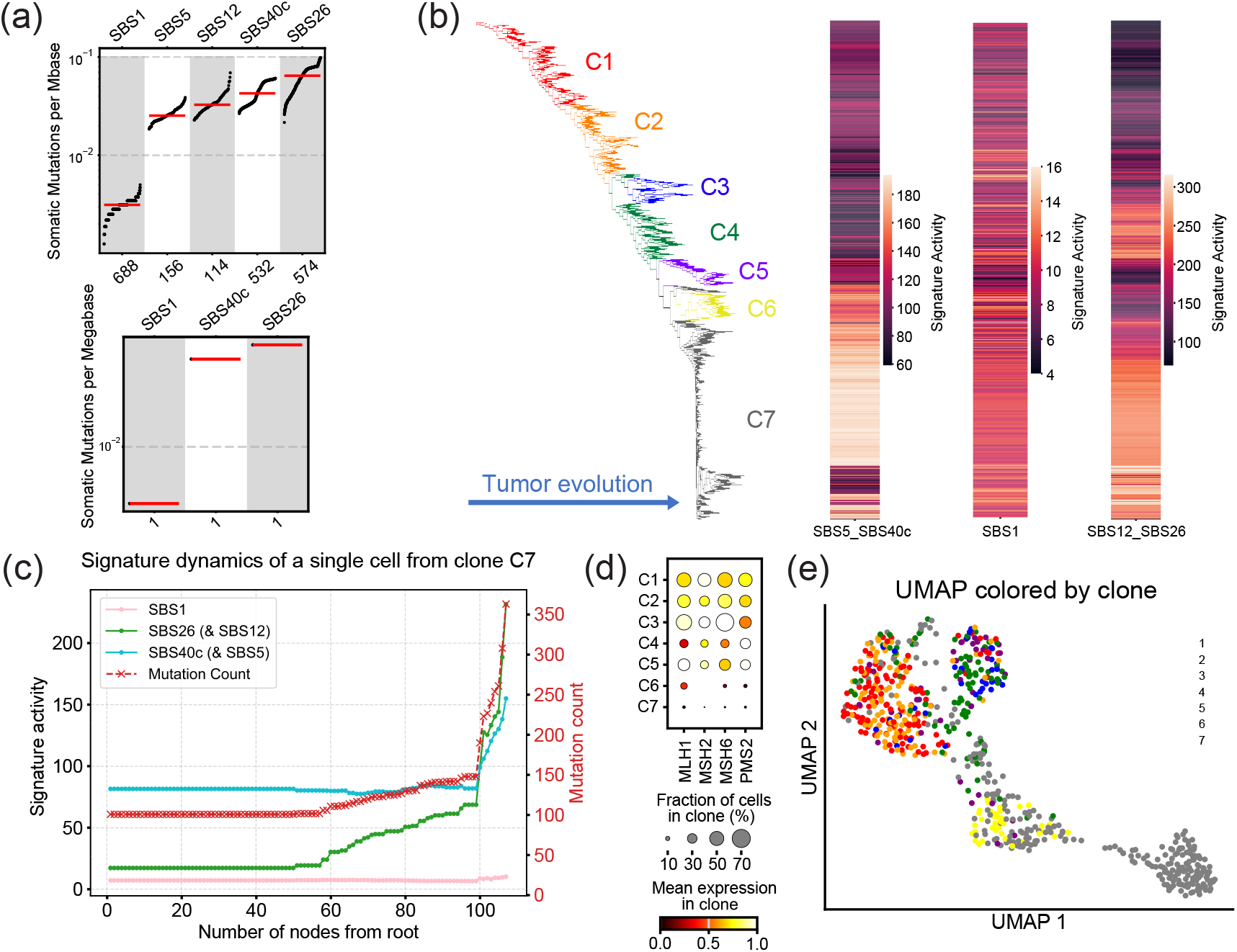
SigDyn mutational signature dynamics and MMR-related gene expression in the breast IDC sample. (a) Mutational signatures at single-cell and pseudo-bulk levels, showing detected COSMIC signatures with respective numbers of samples and activity (mutation counts). (b) Phylogenetic tree inferred from single-cell mutation profiles using FastTree 2, colored by identified clone. Adjacent heatmaps depict absolute activity or exposure of SBS1 and signature pairs SBS12(&26) and SBS5(&40c) across all cells. (c) Dynamics of mutational signatures along the evolutionary trajectory from the root to a randomly chosen cell from clone 7. The left and right axes denote respectively signature exposure and mutation count, while the horizontal axis depicts tree depth, for each node in the trajectory. (d) Dot plot showing the average expression of MMR-related genes (MLH1, MSH2, MSH6, and PMS2) across the seven clones. (e) UMAP embedding plot of single-cell gene expression profiles, colored by clone.

Our results suggest that single-cell sequencing could offer added granularity on mutational processes, especially relevant for heterogeneous samples (Rubanova et al., 2020). However, single-cell RNA-seq analysis has tradeoffs, where lower coverage and transcriptome focus might lead to undetected mutational processes with lower mutation rates. Signature extraction from single-cell DNA-seq could help, albeit at increased cost

### SigDyn on breast invasive ductal carcinoma

We analyzed SigDyn mutational signature dynamics for the breast IDC dataset in greater detail. The independently derived tumor phylogeny and signature activities were visibly aligned, with activities generally increasing along the tumor evolution, most prominently for SBS12/26 and SBS5/40c compared to SBS1 (Fig. 3b,c). The inferred tree clones also showed coherent signature activity (Fig. 3b): clone 4 (green) halfway the evolution tree showed relatively low SBS5/40c and high SBS12/26 activity, whereas clone 7 (grey) farthest in the evolution had relatively high activity for both SBS5/40c and SBS12/26. Additionally, clone 7 contained a subclone with the highest SBS12/26 activity in the IDC sample, and modest activity of SBS5/40c (Fig. 3b,c). Note that we assessed each pair of complementarily detected signatures, SBS5/40c and SBS12/26, together using the sum of activities.

To investigate the relevance of the SigDyn signature dynamics, we looked into the relation between MMR deficiency denoted by SBS26 activity and the transcription levels of four key MMR genes: MLH1, MSH2, MSH6, and PMS2 (Cheng *et al*., 2019). Expression of these MMR genes was associated with the ordering of the cells in the phylogenetic tree (Fig. 3d) and anti-correlated with SBS26 activity (Spearman correlations: MLH1 −0.34, MSH2 −0.31, MSH6 −0.40, PMS2 −0.32; all with *p*-value *<* 0.001), suggesting that MMR function was disrupted over the tumor evolution. Clone 7, near the terminal branches of the tree, showed the highest average SBS26 activity and lowest average expression of the four MMR-related genes (Fig. 3d, Supplementary Fig. S4). This clone also occupied a distinct region in a UMAP projection of the single-cell expression profiles (Fig. 3e), confirming its unique mutational and transcriptional characteristics.

### SigDyn on laryngeal squamous cell carcinoma

To further assess the applicability of SigDyn, we applied it to an LSCC sample comprising 56,145 cells. Following variant calling and filtering based on cohort normals, we identified 730 variant sites. Effects of variant filtering and tree rooting were clearly visible on the inferred tumor phylogeny (Fig. 4a). Variant filtering promoted the organization of single-cells into coherent groupings per region of tissue sampled. Tree rooting produced a natural ordering of the different tissue regions, consistent with the expected tumor evolution: normal tissue (N) near the root, followed by tumor margins (R) and primary tumor (T) clones, with two apparent lymph-node metastasis (L) subclones associated with one of the primary tumor clones. Just like for the IDC dataset, ordering of the LSCC tumor regions in the tree was consistent with organization in the gene expression UMAP space (Supplementary Fig. S5).

**Figure 4.**
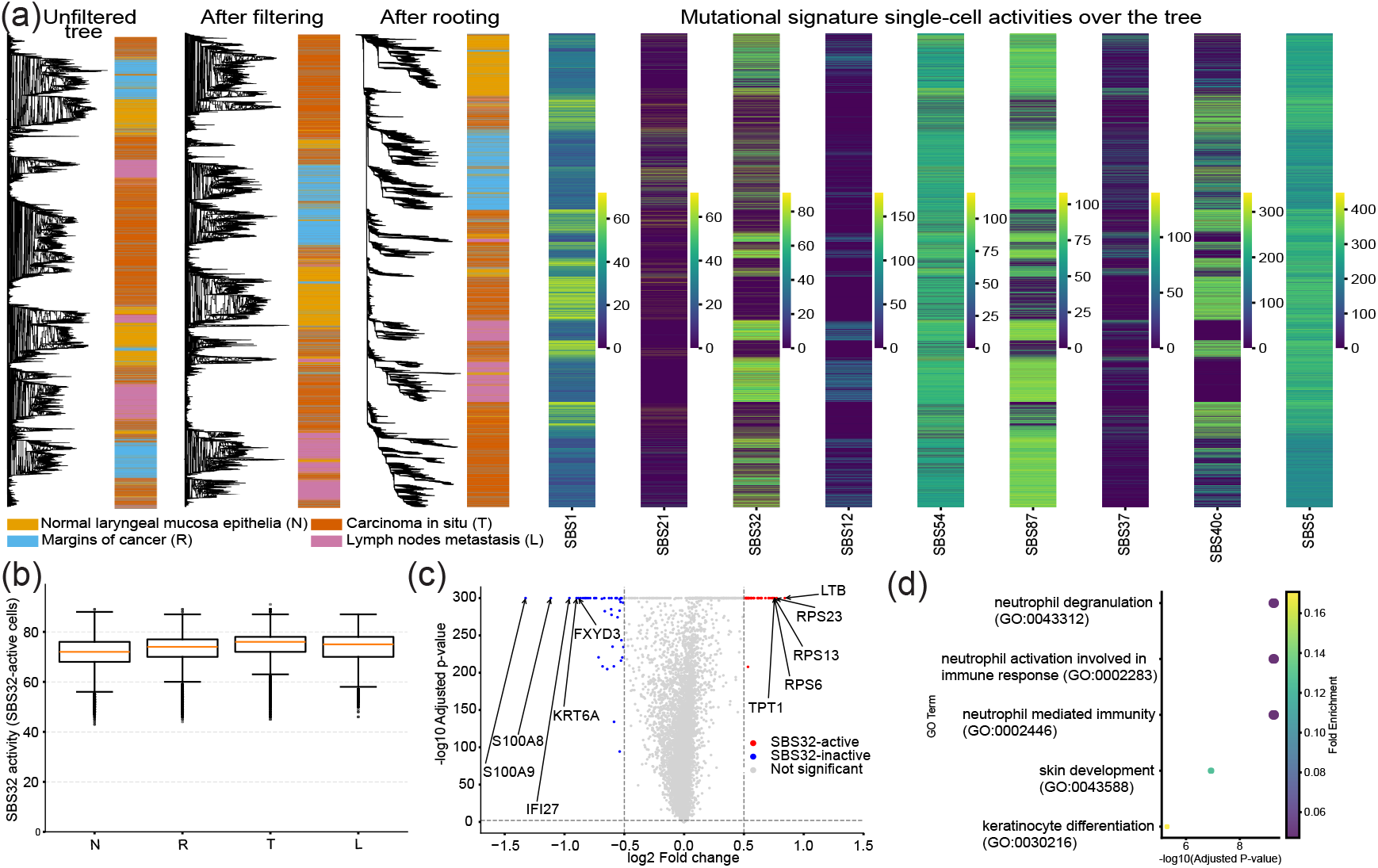
SigDyn mutational signature dynamics and gene expression and functional analysis for the LSCC sample. (a) Phylogenetic trees, annotated by region, inferred: (left) using all unfiltered variants, (middle) using only somatic or filtered variants, (right) using somatic variants and rooting; Heatmaps of single-cell exposures for the 9 mutational signatures present in *>* 10% of the cells. (c) Volcano plot of differentially expressed genes between SBS32-active (red) and SBS32-inactive (blue) cells. (d) Top 5 enriched functional GO terms for genes upregulated in SBS32-inactive cells.

For LSCC, SigDyn identified 4 signatures based on the pseudo-bulk aggregate (Supplementary Fig. S6), namely: SBS54, linked to germline contamination (Alexandrov *et al*., 2020); and clock-like signatures (SBS1, SBS5, SBS40c), which appeared in most cells and generally correlate with age. Overall, these pseudo-bulk signatures could be less relevant to explain the progression of this specific head and neck tumor, reinforcing the need to analyze tumors at single-cell level.

At single-cell level, SigDyn identified 9 mutational signatures present in at least 10% of the cells, including all 4 also detected in the pseudo-bulk (Fig. 4, Supplementary Figs. S7, S8). Mismatch repair deficiency signature SBS21 was detected in 15% of the cells with one of the lowest mutational burdens amongst all signatures (Fig. 4a). Signatures of unknown etiology were identified as well (SBS12, SBS40c, and SBS37). Signature SBS12 was only moderately similar to the identified MMR signature SBS21 (cosine similarity 0.68), suggesting the two were distinct. Moreover, both signatures did not reveal strong associations with MMR-related gene expression (Spearman correlations: for SBS21 MLH1 0.10, MSH2 0.09, MSH6 0.10, PMS2 0.06; for SBS12 MLH1 -0.04, MSH2 -0.02, MSH6 -0.03, PMS2 -0.02; all with *p*-value *<* 0.001). Finally, we identified signatures associated with patient treatments, namely SBS87 with thiopurine chemotherapy and SBS32 with the azathioprine immunosuppressant drug. Treatment history was unavailable, making interpretation of the signatures difficult. However, since SBS32 has been reported in squamous cell carcinomas (SCCs, Inman *et al*. (2018)), we examined it further.

Signature SBS32 was detected in approximately half of all cells (∼26k). The lymph-node metastasis (L) contained the largest proportion of SBS32-active cells, with SBS32 activity *>* 0, amongst all 4 tissue regions (N: 62%, R: 35%, T: 30%, L: 80%). Moreover, the activity in SBS32-active cells was only marginally higher in the primary tumor (T) and metastasis (L) than in the normal tissue (N) (Mann-Whitney U test: L vs N, *p* = 5.9*×*10^−116^, *r* = 0.22; T vs N, *p* = 3.4*×*10^−147^, *r* = 0.25; Fig. 4b). In summary, metastasis was enriched with SBS32-active cells, but these did not show large changes in activity compared to SBS32-active cells in normal tissue. This change in tumor composition suggested that SBS32-active cells could have contributed to tumorigenesis or influenced the metastatic potential of the tumor.

Notably, this was consistent with the fact that signature SBS32 has previously been linked to treatment with azathioprine, an immunosuppressant associated with increased risk of SCCs (Harwood *et al*., 2013). Differential expression analysis further revealed 276 genes with significantly altered expression between SBS32-inactive and SBS32-active cells (Fig. 4c). Follow-up GO term overrepresentation analysis indicated that the 144 genes upregulated in SBS32-inactive cells showed an association with immune response (Fig. 4d), including the top three genes with the largest changes: S100A8, S100A9, and IFI27. The S100A8 and S100A9 genes, members of the S100 protein family, are activated during inflammation and modulate the immune response by promoting leukocyte recruitment and cytokine secretion (Wang *et al*., 2018). The IFI27 gene is involved in regulation of type I and III interferon-mediated responses, RIG-I-like receptor signaling, and innate immune processes in head and neck squamous cell carcinoma (Wang *et al*., 2025). This pattern aligned with the reported etiology of SBS32 and provided additional support for its detection at single-cell level, which was missed in the pseudo-bulk.

## Conclusion

We introduced SigDyn, a framework enabling single-cell analysis of signature dynamics to characterize mutational processes over the evolution of a tumor. Methodologically, SigDyn first detects somatic variants, which are then used to infer a tumor phylogeny and estimate the activity of signatures along such phylogeny.

We acknowledge the challenges associated with identifying variants from single-cell sequencing data, especially focusing on the transcriptome. At the same time, the many cells allow borrowing of information to increase the reliability of variant detection. In addition, SigDyn analysis does not rely on individual variants in isolation, and patterns captured by both phylogeny inference and signature detection are mostly supported by collections of variants.

Notably, SigDyn exposed relevant tumor phylogenies and signature dynamics for breast invasive ductal carcinoma (IDC) and laryngeal squamous cell carcinoma (LSCC) samples.

Ordering of cells along the inferred tumor evolution was well supported by changes in gene expression, including of specific markers like tumor suppressor CST6 in IDC. For LSCC, the tumor phylogeny recapitulated progression from normal to metastasis, in accordance with the sampled tissue regions.

Mutational signature analysis emphasized the potential of single-cell resolution to deliver granular insight into mutational processes. Especially for LSCC, pseudo-bulk analysis identified mostly general age-related signatures, while relevant tumor-specific signatures associated with MMR deficiency or patient treatments were only detected at single-cell level.

Additionally, SigDyn revealed interesting signature dynamics. For IDC, contribution of signature SBS26 increased with tumor evolution, pointing to active acquisition of mutations due to MMR deficiency. For LSCC, the metastasis contained a far larger fraction of SBS32-active cells, whereas SBS32 activity or mutational burden of SBS32-active cells varied marginally across regions from normal tissue to tumor metastasis. This change in cell composition suggested that SBS32-active cells could have influenced tumorigenesis and metastatic potential, consistent with a link of SBS32 to immunosuppressant azathioprine, associated with increased risk of squamous cell carcinoma.

Overall, our results show that SigDyn is designed as a versatile framework that can be used to study tumor heterogeneity, trace subpopulation evolution, and investigate dynamics of mutational processes of individual tumors.

## Data availability

The data underlying this article is publicly available: for IDC, BAM files are at 10xgenomics.com/datasets/; for LSCC, FASTQ files are in the Gene Expression Omnibus (GSE206332).

