## Supplementary Information for "SigDyn: single-cell mutational signature dynamics"

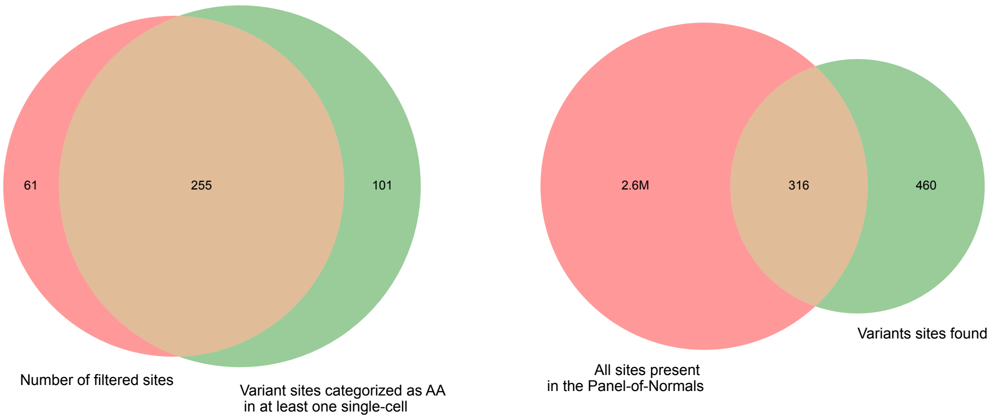


**Figure S1** Comparison of detected variant sites with positions present in the 1000 Genomes Panel-of-Normals (PON). On the left panel, the overlap between filtered positions (PON ∩ dataset) and sites that were assigned an homozygous variant (AA) in at least one single cell. On the right panel, the intersection between all sites present in the PON and all variant positions detected in our dataset. For the right panel only, the areas are shown using log-scaled set sizes.

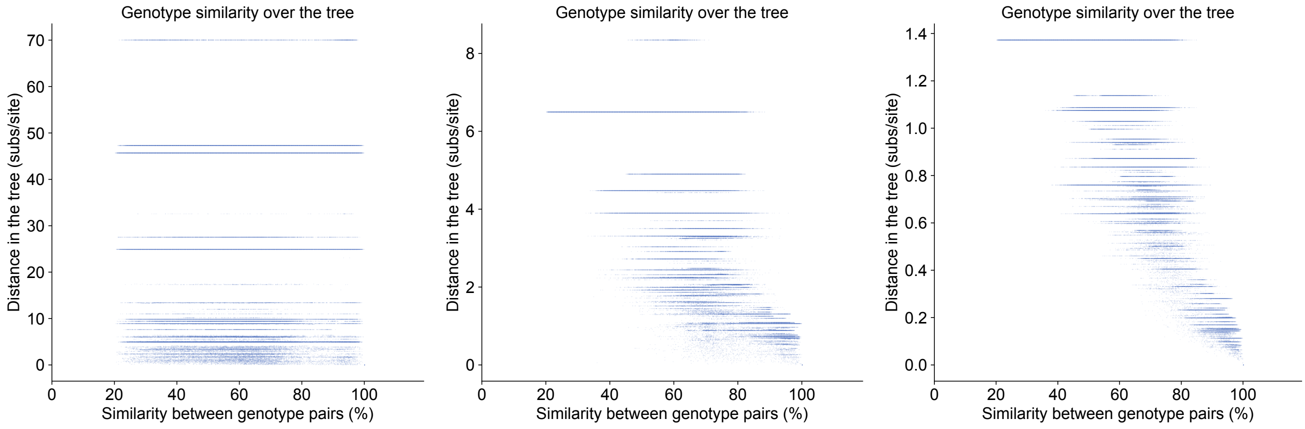


**Figure S2** Relation between single-cell genotype similarity and distance in the tumor phylogenetic tree inferred by SigDyn using BEAST. Panels correspond to (left) 10 thousand iterations; (middle) 1 million iterations; (c) 100 million iterations.

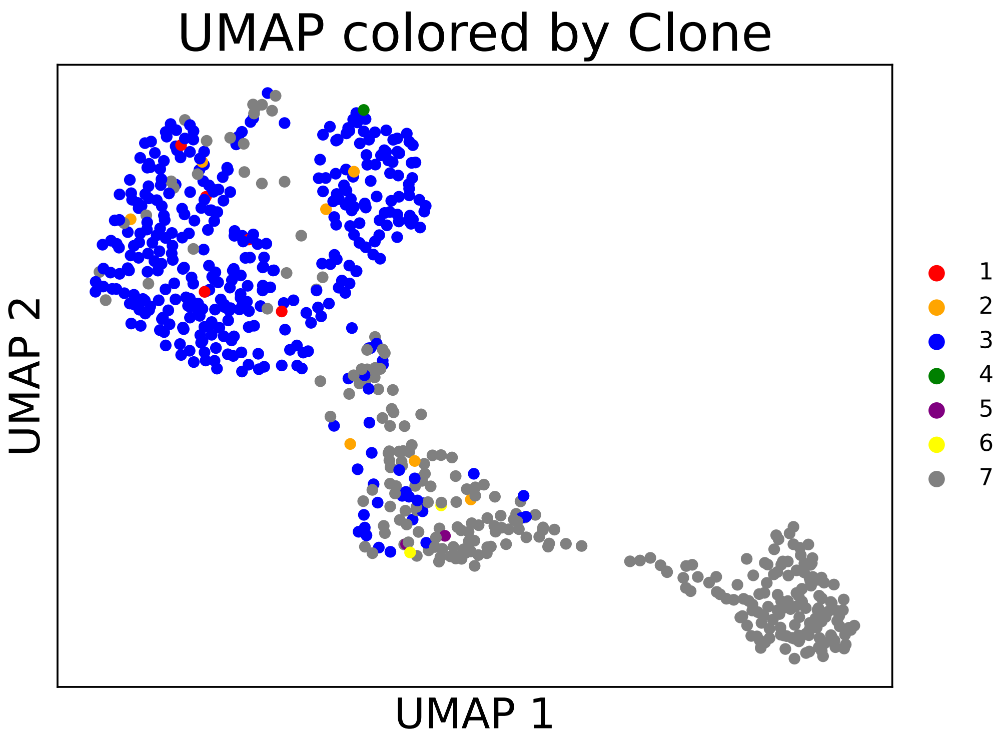


**Figure S3** UMAP embedding plot of single-cell gene expression profiles, colored by BEAST-derived clones.

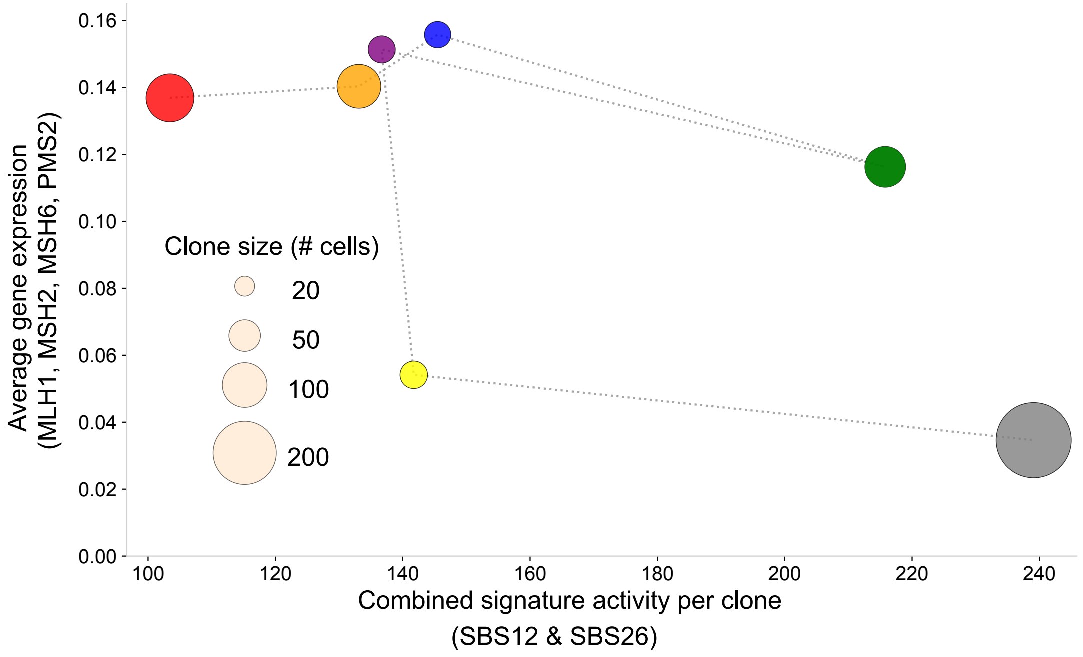


**Figure S4** Relationship between mismatch repair gene expression and mutational signature activity across clones. Each circle represents a clone, plotted by its summed activity of SBS12 and SBS26 (x-axis) and mean expression of mismatch repair genes (MLH1, MSH2, MSH6, PMS2; y-axis). Circle sizes indicate clone sizes (number of cells). Values are averaged per clone. The dotted lines between circles follow the same direction as the clones in the tree.

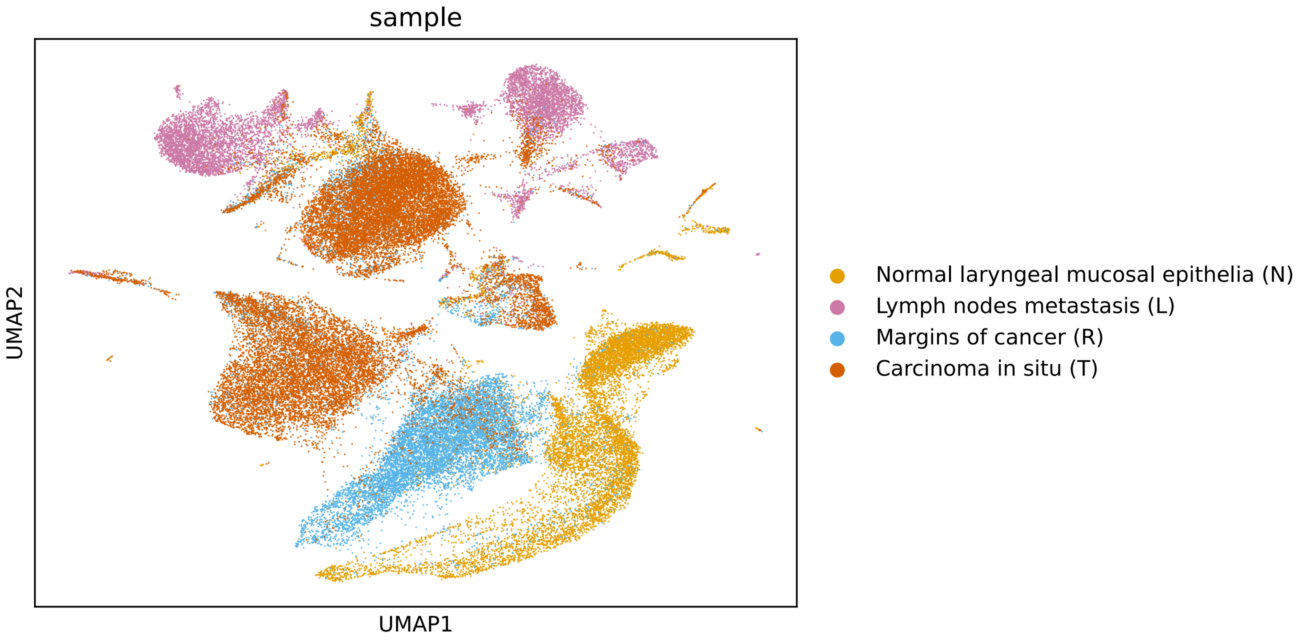


**Figure S5** UMAP projection of four regions of LSCC based on gene expression. Cells are colored by region of origin.

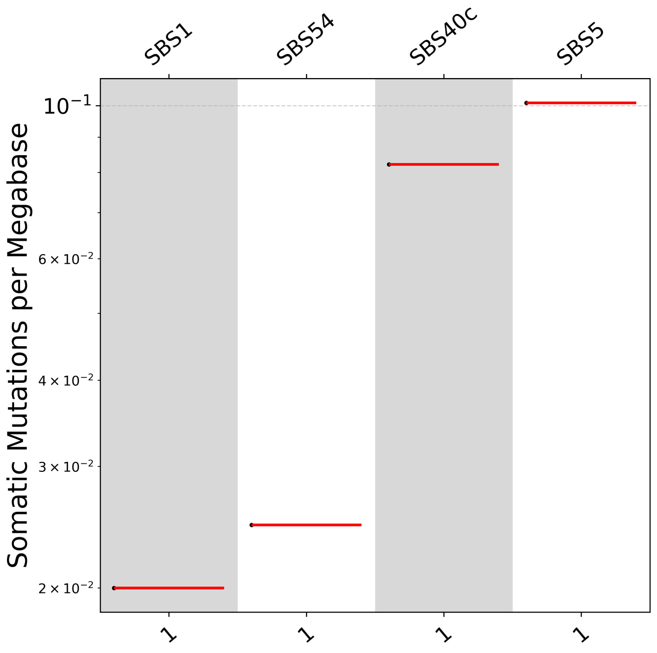


**Figure S6** Assigned mutational signatures at the pseudo-bulk level.

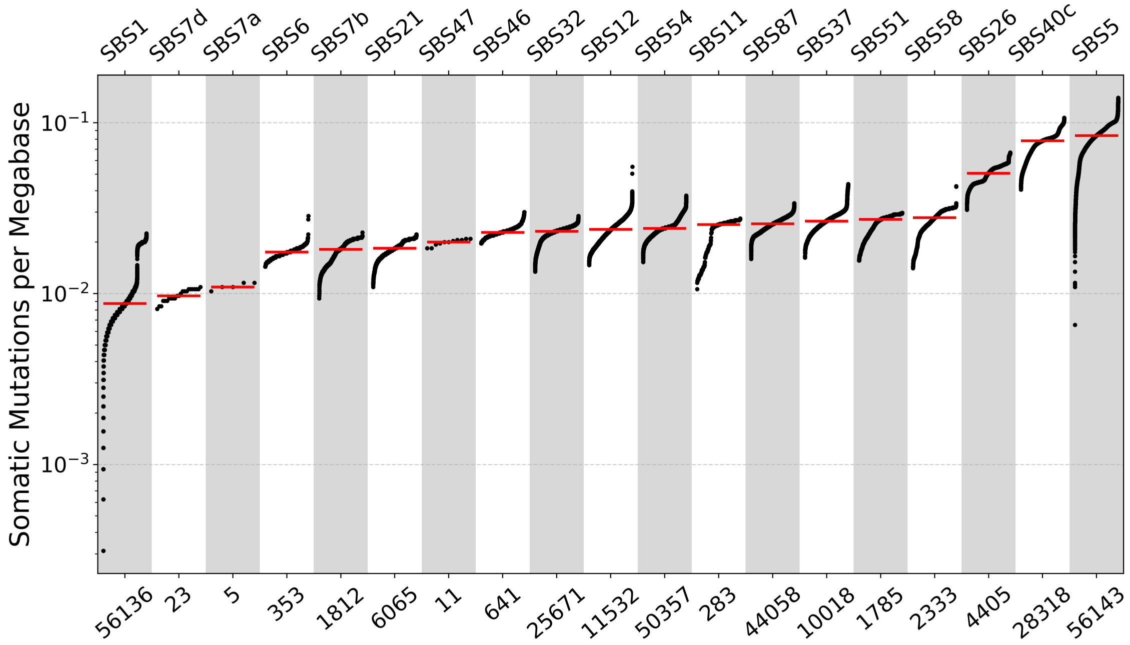


**Figure S7** All initially assigned mutational signatures at the single-cell level.

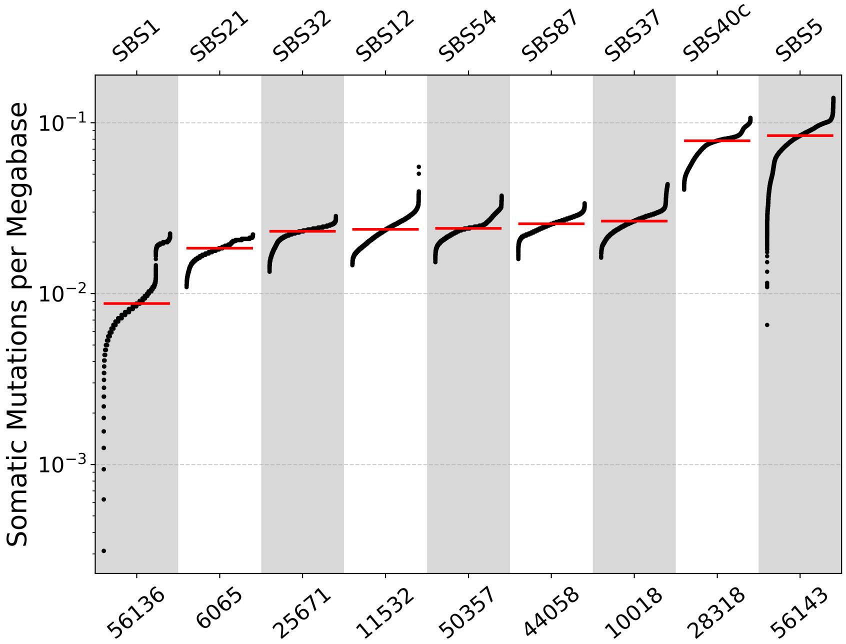


**Figure S8** Assigned mutational signatures at the single-cell level, occurring in more than 10% of cells.
